# Milkweed plants bought at nurseries may expose monarch caterpillars to harmful pesticide residues

**DOI:** 10.1101/2022.03.30.486459

**Authors:** Christopher A. Halsch, Sarah M. Hoyle, Aimee Code, James A. Fordyce, Matthew L. Forister

## Abstract

The decline of monarch butterflies in both the eastern and western United States has garnered widespread public interest. Planting milkweeds, their larval host plants, has been promoted as one action individuals can take, but little is known with respect to potential pesticide contamination of store-bought milkweeds. In this study, we collected leaf samples from 235 milkweed plants purchased at 33 retail nurseries across the US to screen for pesticides. Across all samples, we detected 61 different pesticides with an average of 12.2 (±5.0) compounds per plant. While only 9 of these compounds have been experimentally tested on monarch caterpillars, 38% of samples contained a pesticide above a concentration shown to have a sub-lethal effect for monarchs. We detected only a modest predictive ability of retailer size and milkweed species; and plants with labels advertising their value for wildlife did not have fewer pesticides at concentrations known to have a negative effect on monarchs. These results demonstrate the extensiveness of pesticide exposure within nursery milkweeds and the potential impacts on monarchs and other insects exposed to store-bought plants.

**Highlights:**

- Milkweeds were collected from stores in the United States and screened for pesticides.
- We detected multiple pesticides in every milkweed plant sampled.
- Over one third of samples contained a pesticide at a known harmful concentration for monarchs.
- Plants labeled as wildlife-friendly did not have fewer potentially harmful compounds.

## 1. Introduction

Insect populations are facing an expansive and interacting set of stressors (Habel et al., 2019; Sánchez-Bayo and Wyckhuys, 2019; Wagner et al., 2021). Among native insects declining in the US, the monarch (*Danaus-plexippus*) is a widely recognized butterfly whose declines have been substantial in both the eastern and western populations (Espeset et al., 2016; Pelton et al., 2019; Thogmartin et al., 2017). The causes of these declines are complex and region-specific, but proposed hypotheses implicate climate change, the loss of overwintering habitat, natural enemies, and pesticides, especially herbicides affecting the abundance of native milkweeds (*Asclepias* spp.) (Crone et al., 2019; Zylstra et al., 2021). Many monarch conservation strategies have been suggested, from changes in agricultural practices to the planting of milkweeds by individuals in gardens and yards (Thogmartin et al., 2017). For people wanting to help imperiled insects, planting larval hosts or adult nectar sources is a seemingly simple action, however the practices used to bring nursery plants to shelf often involve pesticide treatments (Krischik et al., 2015; Lentola et al., 2017). A recent study found pesticides in milkweeds growing across diverse landscapes in the Central Valley of California, including up to 31 compounds in individual retail plants (Halsch et al., 2020). Milkweeds in that study were purchased from only two nurseries in one metropolitan area, thus one of the conclusions was that retail locations required further investigation.

In this study, we address the need to better understand contamination of retail plants by quantifying the concentrations of pesticides (insecticides, fungicides, and herbicides) found in the leaves of milkweed plants sold in nurseries across the United States. Retail outlets ranged from local nurseries to large national chains and were all in areas where monarchs breed. First, we present an exploratory analysis of the detected compounds. Second, we examine associations between observed pesticide concentrations and factors relevant to a milkweed buyer. Finally, we offer suggestions for buyers interested in planting milkweeds for monarch conservation.

## 2. Methods

### 2.1 Sample collection

Plants from five milkweed species (*Asclepias curassavica, Asclepias fascicularis, Asclepias incarnata, Asclepias speciosa*, and *Asclepias tuberosa*) were purchased in person from 33 stores across 15 states from May 15 to June 29, 2021. When more than one species could be purchased at a single location, that was done; however, most stores only sold one species (Table S1). Whenever possible, five plants of each species were purchased from each store (fewer plants were available at two instances). These collections were intended to represent a sample of milkweed plants available to the public across the monarch’s migratory range. After purchase, collectors clipped at least 5 grams of leaves from each individual plant, and wrapped samples in tin foil stored in sealed plastic bags. Clippers were cleaned with soap and water or rubbing alcohol between samples. Samples were shipped with ice packs to a central location for temporary storage in a freezer, and ultimately shipped on dry ice to the Cornell Chemical Ecology Core Facility (Cornell University, Ithaca, NY) for chemical analysis. In one instance, 10 leaf samples from two sets of plants were collected immediately after purchase, and another 10 were collected after growing the same individual plants outdoors in pots with daily watering for two weeks (these additional samples were excluded from the primary analyses but are discussed below).

### 2.2 Chemical analysis

Frozen milkweed leaves were extracted by a modified version of the EN 15662 QuEChERS procedure (Standardization, 2008) and screened for 92 pesticides (including some metabolites and breakdown products) by liquid chromatography mass spectrometry (LC-MS/MS). Five grams of frozen leaves were mixed with 10 mL of acetonitrile (five grams was the target weight; samples ranged from 0.88 to 5.09 grams and were prepared accordingly). The leaves were homogenized for 1 min using ceramic beads (2.8 mm diameter) and a Bead Ruptor 24 (OMNI International, United States). After homogenization, 6.5 g of EN 15662 salts (4 g MgSO4; 1 g NaCl; 1 g sodium citrate tribasic dihydrate; 0.5 g sodium citrate dibasic sesquihydrate) were added. Samples were homogenized for an additional 30 sec and centrifuged at 7300 × g for 10 min. One milliliter of supernatant was collected and transferred to a d-SPE (dispersive solid phase extraction) tube containing 150 mg PSA and 900 mg MgSO4. The samples were vortex-mixed for 1 min and centrifuged at 7300 × g for 5 min. Supernatant (294 μL) was collected and six μL of a solution containing three internal standards (0.3 μg/mL 13C6-metalaxyl; 0.3 μg/mL 2H3-pyraclostrobin; 0.15 μg/mL 2H4-fluopyram) was added. The samples were filtered (0.22 μm, PTFE) and stored at −20°C.

Sample analysis was carried out with a Vanquish Flex UHPLC system (Dionex Softron GmbH, Germering, Germany) coupled with a TSQ Quantis mass spectrometer (Thermo Scientific, San Jose, CA, United States). The UHPLC was equipped with an Accuity BEH C18 column (100 mm × 2.1 mm, 1.7 μm particle size, part no. 186002352, Waters, Milford, MA). The mobile phase consisted of (A) water with 2 mM ammonium formate and 0.1% formic acid, and (B) acetonitrile/ water (98:2, v/v) with 2 mM ammonium formate and 0.1% formic acid. The temperature of the column was maintained at 40°C throughout the run and the flow rate was 300 μL/min. The elution program was as follows: 1.5 min prior to injection (2% B, equilibration), 0– 0.5 min (2% B, isocratic), 0.5–15 min (2–70% B, linear gradient), 15–17 min (70–100% B, linear gradient), 17–20 min (100% B, column wash), 20–20.2 min (100–2% B, linear gradient), 20.2–23 min (2% B, re-equilibration). The flow from the LC was directed to the mass spectrometer through a heated electrospray probe (H-ESI). The settings of the H-ESI were as follows: spray voltage 2000 V for both positive and negative mode, sheath gas 55 (arbitrary units), auxiliary gas 25 (arbitrary units), sweep gas 2 (arbitrary units), ion transfer tube temperature 325°C, and vaporizer temperature 350°C.

Tandem mass spectrometry detection was carried out using the selected reaction monitoring (SRM) mode. Two transitions were monitored for each compound: one for quantification and the other for confirmation. The SRM parameters for each individual pesticide are summarized in Supplementary Table S2. The resolution of both Q1 and Q3 was set at 0.7 FWHM, the cycle time was 0.4 s, and the pressure of the collision gas (argon) was set at 2 mTorr.

### 2.3 Statistical analysis

To examine relationships between pesticides and explanatory variables, we modeled pesticide richness (the number of compounds), pesticide diversity, and the number of exceedances of a published lethal or sub-lethal concentration in monarchs. Diversity was represented as the effective number of compounds by taking the exponential of the Shannon diversity index (Jost, 2006). Exceedance was treated as a binary category: either a sample did or did not contain a pesticide at or above a concentration known to exhibit lethal or sub-lethal effects. The predictor variables we explored were the retailer size (single store, 2-100 stores, or > 100 stores), milkweed species, region (eastern or western US), and whether the plant contained a label at the point of purchase with information or a claim about its value for wildlife (e.g. “attracts butterflies”). Pesticide richness, diversity, and the number of exceedances were each modeled using a generalized mixed effects model with a Poisson error structure (log link), a Gaussian error structure, and a binomial error structure (logit link), respectively. The individual store the plants were purchased from was included as a random intercept to account for store-specific effects not encompassed by other explanatory variables. We could not separately evaluate the identity of suppliers because in most cases wholesalers provided plants to a single store. To determine the significance of the terms from each model a Wald chi-squared test was performed on all model outputs.

Random effects models were built in R using the lme4 package (Bates et al., 2020) and variance partitioning was performed using the partR2 and modEvA packages (Barbosa et al., 2021; Stoffel et al., 2021). Wald chi-squared tests were performed using the car package (Fox and Weisberg, 2019). To better understand compositional differences among samples, we used principal coordinates analysis on a Jaccard distance matrix of the presence or absence of each compound using the ape and vegan packages (Oksanen et al., 2019; Paradis et al., 2020). Finally, to investigate the importance of wildlife labels, we used a random forest classifier with the presence of a label as the response and the presence of each compound as predictor variables using the randomForest package (Breiman et al., 2018).

## 3. Results

We detected 61 unique compounds out of a possible 92 in the test panel: 24 insecticides, 26 fungicides, 10 herbicides, and 1 synergist (Fig. 1). All samples contained at least 2 compounds, and some contained as many as 28. The fungicides azoxystrobin, metalaxyl, and difenoconazole were the most common, each found in over 75 percent of samples. None of the samples contained a pesticide exceeding a known LD_50_ of a monarch, however only 9 of the 61 detected compounds have been tested for lethal effects in monarchs. We did detect exceedances of published monarch sub-lethal effects in 89 samples, driven by azoxystrobin and trifloxystrobin, occurring in 17 of 25 locations (Fig. 1, Table S3, Table S4, Table S5).

**Figure 1.**
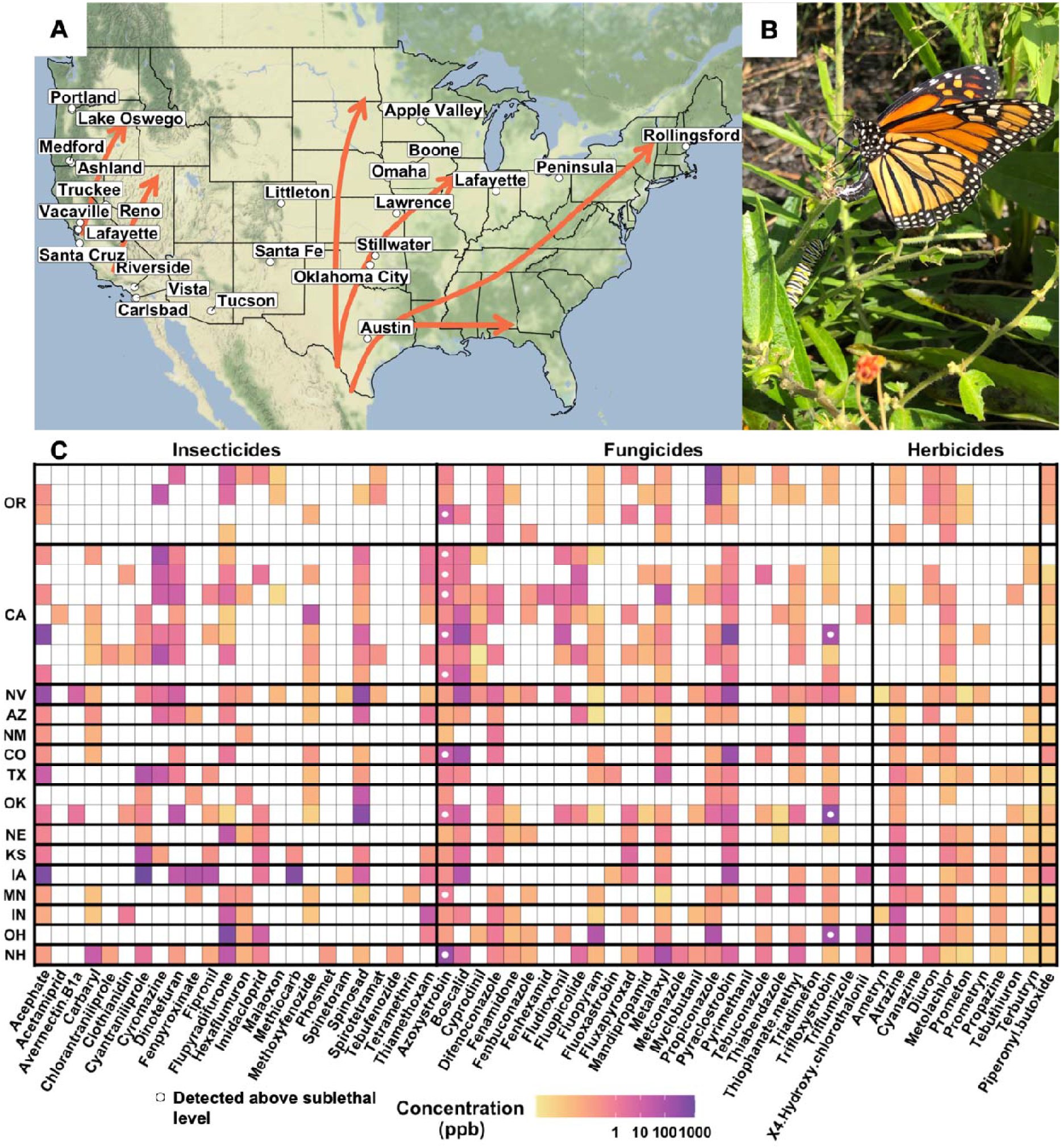
Pesticide and geographic sampling. A) Locations where milkweeds were purchased. Orange arrows denote major monarch migration routes. B) Adult monarch butterfly (*Danaus plexippus*) and caterpillar on *Asclepias tuberosa* (photo credit: MLF). C) Mean concentrations of compounds by city (rows) and by state. Values are shown in parts per billion on a log scale. White circles indicate compounds above a monarch sub-lethal concentration.

Milkweed species and retailer size explained the greatest variation in pesticide richness, and retailer size and the presence of wildlife-friendly labels explained the largest amount of variation in pesticide diversity (Table 1, Fig. S1). Specifically, larger retailers were associated with higher richness (*χ*^2^ = 8.128, *p* = 0.01, Table 2, Table S6, Fig. S2) and higher diversity of compounds (*χ*^2^ = 8.463, *p* = 0.01, Table 2, Table S7, Fig. S3). The presence of a wildlife friendly label was associated with lower diversity due to lower evenness across compounds as we did not find differences in richness (*χ*^2^ = 7.326,*p* < 0.01, Table 2, Table S7, Fig. S3). While pesticide richness and diversity are one aspect of contamination, these metrics are not sufficient for understanding the realized effects of these pesticides on caterpillars. When considering the number of exceedances of a sub-lethal concentration, we found that milkweed species and the presence of wildlife labels explained the most variation (Table 1, Fig. 2). Most notably, plants with a label claiming the plant’s value for wildlife had an increased chance of exceeding a sub-lethal concentration of at least one pesticide (*χ*^2^ = 3.167, *p* = 0.08, Fig. 2, Table S8). Random forest analysis identified azoxystrobin and trifloxystrobin among the most predictive of whether a sample was labeled, both of which were often detected above the sub-lethal threshold in plants labeled valuable for wildlife (Fig. 2). Finally, we did not observe any clustering in response to our predictor variables along the first two PCoA axes and thus patterns of pesticide co-occurrence are apparently not explained by milkweed species, retailer size, region, or labels (Fig. 3).

**Figure 2.**
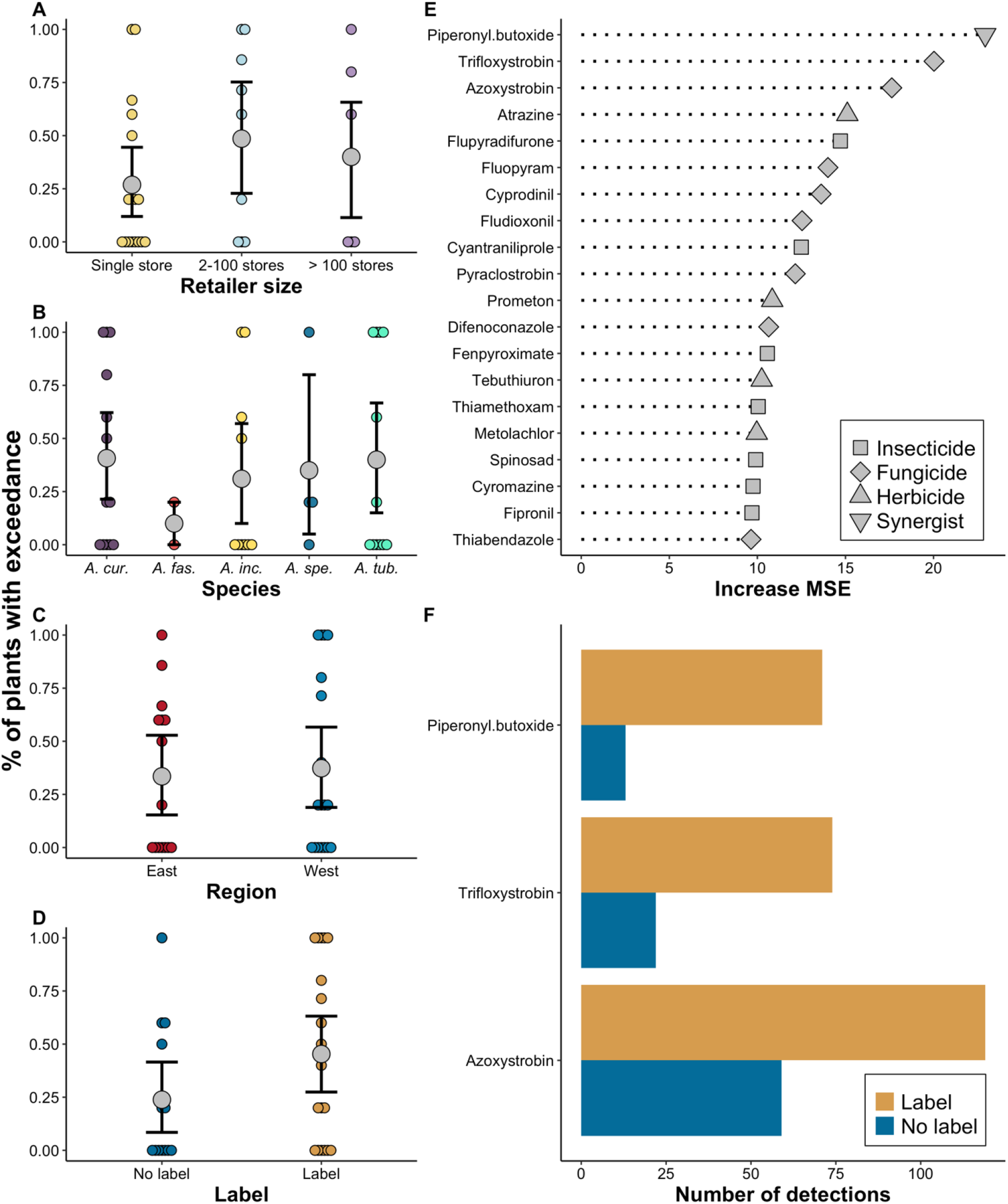
Summary of exceedance associations. Panels A-D show the percentage of plants from each store with at least one exceedance by retailer size, species, region, and wildlife label. Gray points show the mean percentage (bars are standard errors) and points are individual stores. E) Compounds that are the most predictive of whether a sample was labeled as valuable for wildlife based on random forest analysis. F) Differences in the total number of detections between labeled and un-labeled samples for the three most predictive compounds.

**Figure 3.**
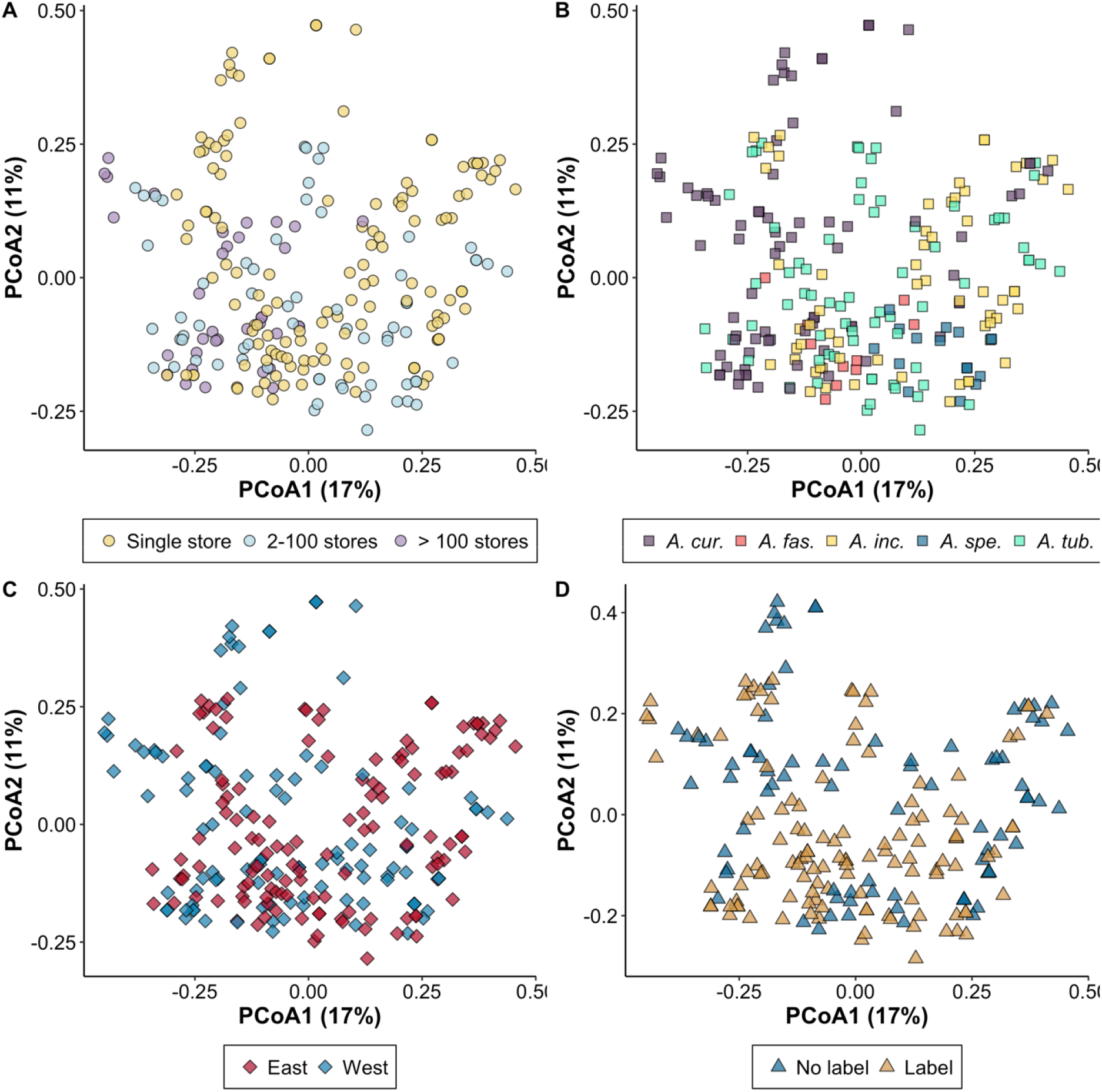
First two axes of a principal coordinates analysis on a Jaccard distance matrix of the presence or absence of each compound by A) retailer size, B) species, C) region, and D) wildlife label.

**Table 1.** Variance explained by covariates predicting pesticide richness, pesticide diversity, and the presence of a lethal or sub-lethal exceedance. The total explained variance is the explained variance of the model when including all terms and the covariate specific rows are the semipartial variance explained by each term.

|  | Pesticide richness | Pesticide diversity | Exceedance |
| --- | --- | --- | --- |
| Total explained variance | 20.5% | 30.0 % | 33.5 % |
| Retailer size | 9.8% | 3.9% | 1.0 % |
| Milkweed species | 9.5 % | 7.9% | 13.2% |
| Region | 0.5% | 0% | 0.0% |
| Label | 0% | 17.2% | 25.4% |

**Table 2.** Means and standard errors of pesticide richness and pesticide diversity for each covariate level.

| Covariate |  | Pesticide richness | Pesticide diversity |
| --- | --- | --- | --- |
| Retailer size | Single store | 11.25 (± 0.39) | 2.98 (± 0.13) |
| | 2-100 stores | 11.85 (± 0.58) | 2.94 (± 0.18) |
| | > 100 stores | 16.43 (± 0.83) | 4.02 (± 0.32) |
| Milkweed species | *A. curassavica* | 12.18 (± 0.59) | 2.69 (± 0.17) |
| | *A. fascicularis* | 13.50 (± 0.67) | 4.53 (± 0.73) |
| | *A. incarnata* | 12.03 (± 0.59) | 3.58 (± 0.22) |
| | *A. speciosa* | 9.12 (± 0.68) | 2.70 (± 0.37) |
| | *A. tuberosa* | 12.92 (± 0.65) | 3.13 (± 0.17) |
| Region | East | 12.32 (± 0.40) | 3.04 (± 0.12) |
| | West | 12.04 (± 0.52) | 3.23 (± 0.18) |
| Label | Yes | 13.33 (± 0.49) | 2.77 (± 0.14) |
| | No | 10.64 (± 0.84) | 3.79 (± 0.20) |

## 4. Discussion

Monarch butterflies are in decline in both the eastern and western United States, leading to an upswell in conservation efforts, including planting milkweeds in gardens and yards. We detected pesticides in milkweeds bought from nurseries in regions of the US where monarchs breed. The number of compounds and their concentrations were highly variable. While we did not detect individual compounds at concentrations known to have lethal impacts on monarchs, the lethal concentrations of most compounds on monarchs are unknown. From the limited set of bioassay experiments that have been performed to test the effects of pesticides, we found that many plants contained pesticides at levels known to have sub-lethal effects. Plants from larger retailers contained higher numbers of pesticides (but not necessarily more sub-lethal exceedances) and, most staggeringly, plants with a label referring to its value for wildlife were almost twice as likely to contain at least one pesticide at a potentially harmful level. This finding is primarily driven by the fungicides azoxystrobin and trifloxystrobin, which were both more common in samples labeled as valuable for wildlife. One possible explanation is that these compounds are not intended to target insects and, thus, might not raise a concern when being used, however, fungicides do have direct impacts on monarchs and other insects (Bernauer et al., 2015; Olaya-Arenas et al., 2020; Su et al., 2014; Tsvetkov et al., 2017; Zhu et al., 2014). Critically, most of the compounds we detected (including some of the most prevalent and those with the highest concentrations) have not been directly tested on monarchs, so harmful levels of contamination are likely underestimated and may be widespread in nursery plants.

Although we did not detect lethal concentrations, we did find extensive potential for sub-lethal effects, which can affect long-term survival and reproduction (Desneux et al., 2006). In monarchs, sub-lethal effects have not often been studied, but a few have been identified.

Olaya□ Arenas et al. (2020) investigated the impacts of six compounds, all of which we detected, and found that larval exposure to azoxystrobin and trifloxystrobin reduce adult monarch wing size (Olaya-Arenas et al., 2020). The concentrations that caused these effects in their study were exceeded on 73 occasions by azoxystrobin and 22 times by trifloxystrobin in our samples. It is noteworthy that similar carryover effects of larval exposure into adult forms have been observed in other non-target butterflies as well (Basley and Goulson, 2018; Whitehorn et al., 2018), further suggesting the potential widespread threat of sub-lethal concentrations on beneficial insects. Other sub-lethal and non-target effects can be realized through pesticides in nectar. Although adult monarchs do nectar at milkweed flowers (and the flowers of many other species), our study was not designed to assay nectar contamination and this remains an important area of future study (Desneux et al., 2006).

While we know little about lethal concentrations and sub-lethal effects, we know even less about how multiple compounds can interact to impact monarchs and other non-target insects. On average, our samples contained 12.2 different pesticides and as many as 28, some of which have been shown to synergistically interact with deleterious effects on the target lepidopteran pest species (Chen et al., 2019; Jones et al., 2012; Liu et al., 2018). In the previously mentioned study, Olaya-Arenas et al. also examined the impacts of a mixture of compounds on monarch caterpillars and did not find strong effects (Olaya-Arenas et al., 2020). These are encouraging findings and a reminder that multiple pesticides do not always result in synergistic negative effects and outcomes can depend on the pesticides present and their concentrations (Cheng et al., 2020; Yu et al., 2016). That said, it is unclear how those results, pertaining to a mixture treatment containing 6 compounds (with 1 insecticide), might apply to our samples which contained, on average, over 12 compounds and often multiple insecticides. Additionally, one hundred of the samples we collected contained piperonyl butoxide, a synergist compound that inhibits a caterpillar’s ability to resist other pesticides through the inhibition of cytochrome P450 enzymes (Young et al., 2005). Short of feeding these specific leaves with complex mixtures to monarch caterpillars, it is likely impossible to predict what the outcomes will be; however, it is difficult to imagine that high pesticide richness will have a positive outcome on monarchs.

These findings raise the possibility for royal concern for the monarch; however, a yet unconsidered and key aspect of these results is the dissipation of compounds over time. Through decomposition, plant growth, wash off, and other factors, the concentration of pesticides are often reduced over time scales of days to weeks, though some compounds may persist much longer (Fantke and Juraske, 2013). Through these processes it is possible that by the time a monarch finds a plant or by the time it completes its development the overall pesticide profile of the leaf will change and become less toxic. In some instances, plants contained a label stating to wait two weeks before feeding to wildlife. To investigate this suggestion, two sets of leaves were purchased from the same store and were screened at two different time points, immediately after purchase and two weeks later, to gain some insight into pesticide dissipation over time. We observed reductions in the concentrations of some pesticides over this time. Compounds like spinosad and acephate which have short half-lives saw the greatest reduction, while other compounds like azoxystrobin (which has a longer half-life) did not change (Table S9, Fig. S4) (Fantke et al., 2014). While it is reassuring that reductions were observed over time in some compounds, azoxytrobin, with the potential for negative effects on monarch caterpillars, did not dissipate in this period. It is also important to note that monarchs are more susceptible to pesticides at earlier instars and even if higher concentration pesticides do dissipate over time, this might be too late for a caterpillar that has hatched onto or has been fed a newly purchased plant (Krishnan et al., 2020). Thus, we can highlight yet another important area for future research: a comprehensive understanding of pesticide persistence in plants sourced from nurseries could lead to important guidelines for private consumers and restoration efforts.

## 5. Conclusions

These findings all lead to the question: what should the average person do if they want to support monarch conservation by planting milkweed? We recommend that plants be purchased from nurseries implementing a robust approach to minimizing their reliance on pesticides, both the total number and concentrations used. This recommendation extends beyond insecticides, as fungicides can also have negative effects on monarch caterpillars and were ubiquitous in our samples. We observed substantial variation in the number of compounds detected per sample, with some samples containing as few as two compounds, indicating that it is possible for nurseries to sell less contaminated plants. Consumers should ask retailers to source plants grown using ecologically sound pest management strategies (Selvaggio and Code, 2020a, 2020b). On the regulatory front, the U.S. Environmental Protection Agency could take steps to address risk, including reducing permissible nursery application rates to prevent residues toxic to pollinators. The monarch has received considerable attention as a declining butterfly in the US, but it is one of many, and may not even be the most dire case (Forister et al., 2021). The threats facing the monarch are the same that are impacting many other native butterflies. The planting of beneficial plants can help some of these imperiled insects, yet, for small scale insect conservation efforts like native plant gardens to be effective, it is critical that nurseries provide plants free from harmful pesticide residues.

## Acknowledgements

We thank all the volunteers and Xerces staff who purchased milkweeds; Wayne Anderson (Cornell Chemical Ecology Core Facility) for performing the chemical analysis; and Sharon Selvaggio for discussion of recommendations.

## Contribution of authors

C.H.: data curation, formal analysis, methodology, visualization, writing – original draft, writing – review and editing; S.H.: data curation, project administration, funding acquisition, writing – review and editing; A.C.: funding acquisition, writing – review and editing; J.F.: methodology, writing – review and editing; M.F.: methodology, supervision, visualization, writing – review and editing.

## Data availability

Data will be made available on Dryad when the paper is accepted for publication.

## Supporting Information

Table S1. Summary of purchased milkweeds.

Table S2. Retention times and optimized SRM acquisition parameters for pesticides and internal standards (RT: Retention time, CE: Collision Energy).

Table S3 Summary of detected compound concentrations in ppb.

Table S4. Sources and thresholds used for pesticides exceedances.

References for Table S4.

Table S5. Number of exceedances of monarch sub-lethal threshold concentrations by compound.

Table S6. Results of Wald chi-aquared test on model predicting pesticide richness.

Table S7. Results of Wald chi-aquared test on model predicting pesticide diversity.

Table S8. Results of Wald chi-aquared test on model predicting pesticide exceedance.

Table S9. Mean change in pesticide concentration across 10 samples over two weeks.

Figure S1. Semi partial variance explained among important predictor variables.

Figure S2. Summary of pesticide richness found in milkweed samples purchased in stores across the United States.

Figure S3. Summary of pesticide diversity found in milkweed samples purchased in stores across the United States.

Figure S4. Changes in pesticide concentrations two weeks after purchase.

