## Supplemental materials for "Milkweed plants bought at nurseries may expose monarch caterpillars to harmful pesticide residues"

### Contents:

Table S1. Summary of purchased milkweeds.

Table S2. Retention times and optimized SRM acquisition parameters for pesticides and internal standards (RT: Retention time, CE: Collision Energy).

Table S3 Summary of detected compound concentrations in ppb.

Table S4. Sources and thresholds used for pesticides exceedances.

References for Table S4.

Table S5. Number of exceedances of monarch sub-lethal threshold concentrations by compound.

Table S6. Results of Wald chi-aquared test on model predicting pesticide richness.

Table S7. Results of Wald chi-aquared test on model predicting pesticide diversity.

Table S8. Results of Wald chi-aquared test on model predicting pesticide exceedance.

Table S9. Mean change in pesticide concentration across 10 samples over two weeks.

Figure S1. Semi partial variance explained among important predictor variables.

Figure S2. Summary of pesticide richness found in milkweed samples purchased in stores across the United States.

Figure S3. Summary of pesticide diversity found in milkweed samples purchased in stores across the United States.

Figure S4. Changes in pesticide concentrations two weeks after purchase.

| Table S1. Summary of purchased milkweeds. | |  |
| --- | --- | --- |
| Milkweed species | Number of plants | Number of retailers |
| *Asclepias curassavica* | 79 | 14 |
| *Asclepias fascicularis* | 10 | 2 |
| *Asclepias incarnata* | 60 | 10 |
| *Asclepias speciosa* | 17 | 4 |
| *Asclepias tuberosa* | 69 | 12 |

| Table S2. Retention times and optimized SRM acquisition parameters for pesticides and internal standards (RT: Retention time, CE: Collision Energy). | | | | | | | | |
| --- | --- | --- | --- | --- | --- | --- | --- | --- |
| Compound | RT (min) | Polarity | Precursor (m/z) | RF Lens (V) | Product 1 (m/z) | CE 1 (V) | Product 2 (m/z) | CE 2 (V) |
| Cyromazine | 1.76 | Positive | 167 | 97 | 85 | 19 | 125 | 18 |
| Acephate | 2.94 | Positive | 184.1 | 66 | 94.9 | 23 | 143 | 10 |
| Dinotefuran | 3.95 | Positive | 203.1 | 67 | 113.1 | 10 | 129 | 12 |
| Thiabendazole | 4.76 | Positive | 202 | 130 | 131 | 33 | 175 | 26 |
| 2,4-DMPF | 5.48 | Positive | 163 | 89 | 107 | 24 | 122 | 16 |
| Thiamethoxam | 5.68 | Positive | 292 | 87 | 181 | 22 | 211.1 | 10 |
| Clothianidin | 6.36 | Positive | 250 | 78 | 131.9 | 17 | 169 | 13 |
| Mevinphos | 6.51 | Positive | 225 | 82 | 127 | 17 | 193 | 10 |
| Imidacloprid | 6.67 | Positive | 256 | 94 | 175 | 18 | 209 | 16 |
| Acetamiprid | 7.17 | Positive | 223 | 92 | 90 | 34 | 125.9 | 22 |
| Flupyradifurone | 7.79 | Positive | 288.9 | 110 | 89.9 | 38 | 126 | 19 |
| Thiacloprid | 8.11 | Positive | 253 | 113 | 90 | 36 | 125.9 | 21 |
| Sulfoxaflor | 8.3 | Negative | 275.9 | 115 | 213 | 17 | 261 | 13 |
| Tebuthiuron | 8.29 | Positive | 229 | 105 | 116 | 27 | 172 | 18 |
| Prometon | 8.43 | Positive | 226.2 | 118 | 142.1 | 23 | 184.1 | 19 |
| Cyanazine | 9.02 | Positive | 241.1 | 124 | 104 | 29 | 214 | 17 |
| Thiophanate-methyl | 9.44 | Positive | 343 | 119 | 151 | 20 | 311 | 10 |
| Ametryn | 9.56 | Positive | 228.1 | 128 | 96 | 26 | 186 | 18 |
| Methoprotryne | 9.63 | Positive | 272.1 | 139 | 170 | 29 | 198 | 23 |
| Malaoxon | 9.74 | Positive | 315 | 101 | 98.9 | 23 | 127 | 12 |
| Bendiocarb | 9.76 | Positive | 224.1 | 81 | 109 | 18 | 167 | 10 |
| Carbofuran | 9.78 | Positive | 222.1 | 79 | 123 | 22 | 165 | 12 |
| 4-Hydroxy-chlorothalonil | 10.12 | Negative | 244.8 | 141 | 174.9 | 27 | 181.9 | 29 |
| Pyrimethanil | 10.23 | Positive | 200.1 | 150 | 82 | 26 | 107 | 24 |
| Carbaryl | 10.26 | Positive | 202 | 73 | 127 | 29 | 145 | 10 |
| Fluometuron | 10.29 | Positive | 233 | 116 | 46 | 18 | 72 | 19 |
| Atrazine | 10.3 | Positive | 216 | 129 | 104 | 29 | 174 | 18 |
| Sulfentrazone | 10.44 | Negative | 384.9 | 200 | 199 | 36 | 307 | 23 |
| Diuron | 10.72 | Positive | 233.1 | 95 | 46 | 17 | 72 | 18 |
| ^13^C_6_-Metalaxyl | 10.75 | Positive | 286 | 90 | 166 | 24 | 226.1 | 18 |
| Metalaxyl | 10.76 | Positive | 280.1 | 104 | 160 | 24 | 220.1 | 14 |
| Cyantraniliprole | 10.84 | Positive | 474.9 | 137 | 285.8 | 14 | 443.9 | 19 |
| Metobromuron | 11.05 | Positive | 258.9 | 99 | 148 | 16 | 169.9 | 19 |
| Prometryn | 11.14 | Positive | 242.1 | 123 | 158 | 23 | 200 | 18 |
| Terbutryn | 11.29 | Positive | 242.1 | 123 | 68 | 40 | 186 | 18 |
| Metazachlor | 11.36 | Positive | 278.1 | 84 | 134.1 | 22 | 210 | 10 |
| Clomazone | 11.65 | Positive | 240.1 | 89 | 89 | 47 | 124.9 | 21 |
| Propazine | 11.8 | Positive | 230.1 | 130 | 146.1 | 23 | 188.1 | 18 |
| Chlorantraniliprole | 11.8 | Positive | 483.9 | 137 | 285.8 | 11 | 452.9 | 17 |
| Methiocarb | 12.27 | Positive | 226.1 | 84 | 121 | 19 | 169 | 10 |
| Phenmedipham | 12.32 | Positive | 301 | 108 | 136 | 20 | 168 | 10 |
| Fluazifop | 12.41 | Positive | 328 | 144 | 254 | 27 | 282 | 19 |
| Flumioxazin | 12.5 | Positive | 355 | 191 | 299 | 29 | 327 | 21 |
| Fludioxonil | 12.62 | Negative | 246.9 | 130 | 126 | 32 | 180 | 30 |
| Spirotetramat | 12.65 | Positive | 374.1 | 140 | 216.1 | 34 | 302.1 | 17 |
| Cyprodynil | 12.66 | Positive | 226.1 | 136 | 93 | 35 | 108.1 | 27 |
| Bromuconazole | 12.7 | Positive | 377.9 | 151 | 159 | 29 | 160.9 | 30 |
| Phosmet | 12.72 | Positive | 318 | 95 | 133 | 36 | 159.9 | 13 |
| Azoxystrobin | 12.96 | Positive | 404 | 127 | 329 | 31 | 372 | 15 |
| Myclobutanil | 13.02 | Positive | 289 | 134 | 70 | 20 | 125 | 34 |
| Fenamidone | 13.03 | Positive | 312 | 112 | 92 | 24 | 236.1 | 14 |
| Triadimefon | 13.11 | Positive | 294 | 111 | 69 | 21 | 197 | 16 |
| Fluxapyroxad | 13.17 | Positive | 382 | 133 | 342 | 21 | 362 | 15 |
| Boscalid | 13.19 | Positive | 342.9 | 125 | 139.9 | 18 | 307 | 20 |
| Mandipropamid | 13.24 | Positive | 412.1 | 126 | 328 | 15 | 356 | 10 |
| Fenhexamid | 13.26 | Positive | 302 | 136 | 55 | 34 | 97 | 24 |
| Spinosad | 13.74 | Positive | 732.4 | 168 | 98 | 43 | 142.1 | 29 |
| Napropamide | 13.39 | Positive | 272.1 | 104 | 171 | 19 | 199 | 13 |
| Tetraconazole | 13.44 | Positive | 372 | 152 | 70 | 23 | 159 | 31 |
| Fluopicolide | 13.45 | Positive | 382.9 | 131 | 144.9 | 48 | 172.8 | 24 |
| Tebuconazole | 13.49 | Positive | 308.1 | 130 | 70 | 23 | 124.9 | 37 |
| d_4_-Fluopyram | 13.53 | Positive | 401 | 139 | 177 | 28 | 208 | 21 |
| Fluopyram | 13.55 | Positive | 397 | 126 | 173 | 29 | 208 | 22 |
| Methoxyfenozide | 13.61 | Positive | 369.1 | 77 | 149 | 17 | 313.1 | 10 |
| Diflubenzuron | 13.71 | Positive | 311 | 100 | 141 | 32 | 158 | 13 |
| Fenbuconazole | 13.81 | Positive | 337 | 144 | 70 | 21 | 125 | 31 |
| Metolachlor | 13.88 | Positive | 284.1 | 102 | 176.1 | 26 | 252 | 15 |
| Metconazole | 14 | Positive | 320.1 | 131 | 70 | 24 | 125 | 39 |
| Flufenacet | 14.11 | Positive | 364 | 95 | 152 | 19 | 194.1 | 10 |
| Propiconazole | 14.22 | Positive | 342 | 150 | 69.1 | 20 | 158.9 | 30 |
| Dimoxystrobin | 14.17 | Positive | 327.1 | 89 | 116 | 22 | 205.1 | 10 |
| Fumagillin | 14.17 | Positive | 459.1 | 124 | 131 | 26 | 177 | 14 |
| Fluoxastrobin | 14.18 | Positive | 459 | 172 | 188 | 35 | 427 | 18 |
| Tebufenozide | 14.51 | Positive | 353.1 | 82 | 133 | 19 | 297.1 | 10 |
| Spinetoram | 14.91 | Positive | 748.3 | 217 | 98 | 44 | 142.1 | 30 |
| Triflumizole | 14.85 | Positive | 346 | 90 | 73.1 | 16 | 278.1 | 10 |
| Difenoconazole | 14.95 | Positive | 406 | 165 | 251 | 26 | 337 | 18 |
| Fipronil | 14.85 | Negative | 434.8 | 149 | 249.9 | 27 | 330 | 16 |
| Picoxystrobin | 14.97 | Positive | 368 | 83 | 145 | 21 | 205 | 10 |
| Penthiopyrad | 15.02 | Positive | 360.1 | 126 | 256 | 21 | 276 | 15 |
| d_3_-Pyraclostrobin | 15.44 | Positive | 391 | 105 | 167 | 17 | 197 | 10 |
| Coumaphos | 15.46 | Positive | 363 | 127 | 227 | 26 | 306.9 | 18 |
| Pyraclostrobin | 15.47 | Positive | 388.1 | 114 | 163 | 24 | 194 | 13 |
| Thiobencarb | 15.56 | Positive | 258 | 88 | 89 | 48 | 125 | 20 |
| Hexaflumuron | 15.79 | Negative | 458.7 | 118 | 275.9 | 20 | 439 | 10 |
| Profenofos | 16.21 | Positive | 374.9 | 130 | 304.8 | 19 | 346.8 | 13 |
| Cyflufenamid | 16.21 | Positive | 413.1 | 128 | 241 | 23 | 295 | 15 |
| Indoxacarb | 16.27 | Positive | 527.9 | 164 | 150 | 24 | 203 | 39 |
| Trifloxystrobin | 16.36 | Positive | 409 | 128 | 145 | 44 | 186 | 18 |
| Piperonyl butoxide | 16.73 | Positive | 356.2 | 92 | 119 | 34 | 177 | 12 |
| Tetramethrin | 16.86 | Positive | 332.1 | 106 | 135.1 | 18 | 164 | 24 |
| Fluazinam | 16.95 | Negative | 462.8 | 138 | 397.9 | 16 | 415.9 | 20 |
| Chlorpyrifos | 17.21 | Positive | 349.9 | 105 | 96.9 | 31 | 197.9 | 20 |
| Fenpyroximate | 17.32 | Positive | 422.1 | 146 | 214.1 | 30 | 366 | 16 |
| Avermectin B1a | 17.79 | Positive | 890.3 | 176 | 305.2 | 24 | 567.3 | 13 |

| Table S3. Summary of detected compound concentrations in ppb. We present the mean, median, minimum, and maximum concentrations across all samples (excluding samples where it was not detected) and the number of samples in which each compound was detected. | | | | | |
| --- | --- | --- | --- | --- | --- |
| Compound | Mean | Median | Min | Max | n |
| Acephate | 174.87 | 0.40 | 0.39 | 2197.80 | 89 |
| Acetamiprid | 0.25 | 0.25 | 0.25 | 0.25 | 1 |
| Ametryn | 0.01 | 0.01 | 0.01 | 0.01 | 5 |
| Atrazine | 2.24 | 0.35 | 0.03 | 24.38 | 145 |
| Avermectin B1a | 15.03 | 6.70 | 6.57 | 35.21 | 8 |
| Azoxystrobin | 21.88 | 0.30 | 0.02 | 800.00 | 207 |
| Boscalid | 55.46 | 6.54 | 1.97 | 478.47 | 83 |
| Carbaryl | 9.36 | 0.07 | 0.07 | 173.55 | 34 |
| Chlorantraniliprole | 0.53 | 0.40 | 0.39 | 1.32 | 7 |
| Clothianidin | 1.01 | 0.40 | 0.39 | 2.58 | 14 |
| Cyanazine | 0.36 | 0.21 | 0.20 | 1.14 | 12 |
| Cyantraniliprole | 330.38 | 2.80 | 0.39 | 2197.80 | 54 |
| Cyprodinil | 0.24 | 0.14 | 0.01 | 0.97 | 28 |
| Cyromazine | 57.14 | 19.81 | 0.40 | 307.62 | 46 |
| Difenoconazole | 0.98 | 0.60 | 0.10 | 14.57 | 177 |
| Dinotefuran | 18.84 | 2.93 | 0.39 | 135.74 | 80 |
| Diuron | 0.73 | 0.40 | 0.39 | 2.19 | 38 |
| Fenamidone | 0.15 | 0.13 | 0.13 | 0.41 | 17 |
| Fenbuconazole | 0.54 | 0.40 | 0.39 | 1.63 | 14 |
| Fenhexamid | 13.35 | 13.36 | 13.31 | 13.39 | 3 |
| Fenpyroximate | 33.10 | 0.65 | 0.13 | 156.20 | 21 |
| Fipronil | 20.20 | 0.78 | 0.20 | 249.27 | 28 |
| Fludioxonil | 5.61 | 3.25 | 0.65 | 38.86 | 35 |
| Fluopicolide | 6.62 | 6.39 | 0.20 | 20.71 | 40 |
| Fluopyram | 10.68 | 0.03 | 0.01 | 222.67 | 65 |
| Fluoxastrobin | 0.19 | 0.10 | 0.07 | 0.43 | 7 |
| Flupyradifurone | 115.49 | 2.86 | 0.02 | 1976.28 | 93 |
| Fluxapyroxad | 2.59 | 0.90 | 0.39 | 21.68 | 28 |
| Hexaflumuron | 0.53 | 0.40 | 0.39 | 1.26 | 13 |
| Imidacloprid | 4.50 | 1.60 | 0.39 | 37.69 | 38 |
| Malaoxon | 0.03 | 0.02 | 0.02 | 0.09 | 12 |
| Mandipropamid | 0.40 | 0.07 | 0.07 | 9.11 | 33 |
| Metalaxyl | 13.62 | 0.14 | 0.01 | 744.97 | 178 |
| Metconazole | 3.00 | 3.20 | 0.40 | 6.02 | 5 |
| Methiocarb | 689.61 | 8.10 | 1.27 | 2078.14 | 11 |
| Methoxyfenozide | 4.56 | 0.07 | 0.07 | 65.37 | 41 |
| Metolachlor | 0.20 | 0.17 | 0.04 | 1.20 | 151 |
| Myclobutanil | 0.24 | 0.13 | 0.13 | 1.63 | 14 |
| Phosmet | 2.02 | 2.02 | 2.02 | 2.02 | 1 |
| Piperonyl butoxide | 0.17 | 0.03 | 0.01 | 3.08 | 100 |
| Prometon | 0.03 | 0.01 | 0.01 | 0.13 | 87 |
| Prometryn | 0.04 | 0.02 | 0.02 | 0.11 | 10 |
| Propazine | 0.09 | 0.07 | 0.07 | 0.24 | 38 |
| Propiconazole | 52.76 | 0.24 | 0.20 | 2000.00 | 99 |
| Pyraclostrobin | 70.55 | 0.76 | 0.07 | 1408.20 | 142 |
| Pyrimethanil | 0.07 | 0.07 | 0.07 | 0.07 | 1 |
| Spinetoram | 0.08 | 0.08 | 0.07 | 0.10 | 3 |
| Spinosad | 142.61 | 1.71 | 0.07 | 2109.70 | 101 |
| Spirotetramat | 0.56 | 0.32 | 0.13 | 1.56 | 10 |
| Tebuconazole | 1.46 | 0.67 | 0.65 | 3.74 | 26 |
| Tebufenozide | 0.37 | 0.41 | 0.07 | 0.63 | 3 |
| Tebuthiuron | 0.07 | 0.02 | 0.02 | 0.43 | 23 |
| Terbutryn | 0.03 | 0.02 | 0.02 | 0.19 | 36 |
| Tetramethrin | 0.40 | 0.40 | 0.40 | 0.40 | 1 |
| Thiabendazole | 0.26 | 0.26 | 0.04 | 1.04 | 23 |
| Thiamethoxam | 4.06 | 1.27 | 0.13 | 42.89 | 71 |
| Thiophanate methyl | 1.80 | 0.22 | 0.13 | 29.73 | 78 |
| Triadimefon | 0.86 | 0.40 | 0.40 | 1.87 | 5 |
| Trifloxystrobin | 88.60 | 0.14 | 0.02 | 2016.13 | 116 |
| Triflumizole | 0.07 | 0.04 | 0.04 | 0.17 | 5 |
| X4 Hydroxy chlorothalonil | 57.75 | 71.15 | 1.98 | 188.03 | 11 |

| Table S4. Sources and thresholds used for pesticides exceedances. Krishnan et al. [1,2] reported LC_10,_ LC_50,_ and LC_90_ values from which we used LC_10_ and LC_50_ values. | | | | | |
| --- | --- | --- | --- | --- | --- |
| Study | Compound | Concentration  (ppb) | Effect type | Exposure type | Instar |
| Bargar et al. [3] | Clothianidin | 47 | LC_50_ | Chronic | All instars |
| Krishnan et al. [1] | Chlorantraniliprole | 8.3 | LC_50_ | Acute | 2nd |
| Krishnan et al. [1] | Imidacloprid | 5100 | LC_50_ | Acute | 2nd |
| Krishnan et al. [1] | Thiamethoxam | 3400 | LC_50_ | Acute | 2nd |
| Krishnan et al. [1] | Chlorantraniliprole | 0.49 | LC_10_ | Acute | 2nd |
| Krishnan et al. [1] | Imidacloprid | 1400 | LC_10_ | Acute | 2nd |
| Krishnan et al. [1] | Thiamethoxam | 1400 | LC_10_ | Acute | 2nd |
| Krishnan et al. [2] | Chlorantraniliprole | 1.6 | LC_50_ | Chronic | 2nd – 5th |
| Krishnan et al. [2] | Clothianidin | 740 | LC_50_ | Chronic | 2nd – 5th |
| Krishnan et al. [2] | Imidacloprid | 130 | LC_50_ | Chronic | 2nd – 5th |
| Krishnan et al. [2] | Thiamethoxam | 940 | LC_50_ | Chronic | 2nd – 5th |
| Krishnan et al. [2] | Chlorantraniliprole | 0.38 | LC_10_ | Chronic | 2nd – 5th |
| Krishnan et al. [2] | Clothianidin | 46 | LC_10_ | Chronic | 2nd – 5th |
| Krishnan et al. [2] | Imidacloprid | 36 | LC_10_ | Chronic | 2nd – 5th |
| Krishnan et al. [2] | Thiamethoxam | 420 | LC_10_ | Chronic | 2nd – 5th |
| Olaya‑Arenas et al. [4] | Clothianidin | 15.28 | No effect | Chronic | All instars |
| Olaya‑Arenas et al. [4] | Atrazine | 8.59 | No effect | Chronic | All instars |
| Olaya‑Arenas et al. [4] | S-Metolachlor | 1.23 | No effect | Chronic | All instars |
| Olaya‑Arenas et al. [4] | Azoxystrobin | 0.67 | Sublethal | Chronic | All instars |
| Olaya‑Arenas et al. [4] | Pyraclostrobin | 8.51 | No effect | Chronic | All instars |
| Olaya‑Arenas et al. [4] | Trifoxystrobin | 4.48 | Sublethal | Chronic | All instars |

| Table S5. Number of plant samples with exceedances of monarch sub-lethal threshold concentrations by compound. | |
| --- | --- |
| Compound Name | Number of Exceedances |
| Azoxystrobin | 73 |
| Trifloxystrobin | 22 |

| Table S6. Results of Wald chi-squared test on model predicting pesticide richness. | | | |
| --- | --- | --- | --- |
| Term | $\chi$^2^ | DF | p-value |
| Species | 17.3736 | 4 | 0.0016 |
| Retailer size | 8.1283 | 2 | 0.0172 |
| Region | 0.2970 | 1 | 0.5858 |
| Wildlife label | 0.2474 | 1 | 0.6189 |

| Table S7. Results of Wald chi-squared test on model predicting pesticide diversity. | | | |
| --- | --- | --- | --- |
| Term | $\chi$^2^ | DF | p-value |
| Species | 62.8974 | 4 | < 0.0001 |
| Retailer size | 8.4634 | 2 | 0.0145 |
| Region | 0.0015 | 1 | 0.9694 |
| Wildlife label | 7.3259 | 1 | 0.0068 |

| Table S8. Results of Wald chi-squared test on model predicting pesticide exceedance. | | | |
| --- | --- | --- | --- |
| Term | $\chi$^2^ | DF | p-value |
| Species | 3.5723 | 4 | 0.4670 |
| Retailer size | 1.4082 | 2 | 0.4946 |
| Region | 0.0122 | 1 | 0.9121 |
| Wildlife label | 3.1673 | 1 | 0.0751 |

| Table S9. Mean change in pesticide concentration across 10 samples over two weeks. Rows are shown in ascending order with highest declines at the top. Also see Fig. S4. | |
| --- | --- |
| Compound | Change in concentration (log(ppb + 1)) |
| Spinosad | -0.9649273 |
| Acephate | -0.8258331 |
| Avermectin B1a | -0.6980482 |
| Pyraclostrobin | -0.575131 |
| Propiconazole | -0.5226526 |
| Difenoconazole | -0.4854371 |
| Fludioxonil | -0.4479201 |
| Thiabendazole | -0.3223651 |
| Trifloxystrobin | -0.2748731 |
| Cyantraniliprole | -0.2245888 |
| Fenbuconazole | -0.1976401 |
| Triadimefon | -0.1865242 |
| Dinotefuran | -0.1120412 |
| Thiophanate methyl | -0.1016475 |
| Cyromazine | -0.0931322 |
| Cyprodinil | -0.0743814 |
| Hexaflumuron | -0.0672944 |
| Myclobutanil | -0.0611088 |
| Boscalid | -0.0508832 |
| Piperonyl.butoxide | -0.0438334 |
| Mandipropamid | -0.0338293 |
| Diuron | -0.0336472 |
| Triflumizole | -0.0313887 |
| Flupyradifurone | -0.030495 |
| Carbaryl | -0.0270635 |
| Fenamidone | -0.0244435 |
| Fluxapyroxad | -0.0241914 |
| Atrazine | -0.0202908 |
| Prometryn | -0.0173042 |
| Malaoxon | -0.0165388 |
| Spirotetramat | -0.0122218 |
| Thiamethoxam | -0.0122218 |
| Spinetoram | -0.0067659 |
| Azoxystrobin | -0.0040197 |
| Ametryn | -0.000995 |
| Fluopyram | -0.000995 |
| Prometon | -0.000995 |
| Metolachlor | 0.0151135 |
| Methoxyfenozide | 0.04736105 |
| Metalaxyl | 0.0852799 |

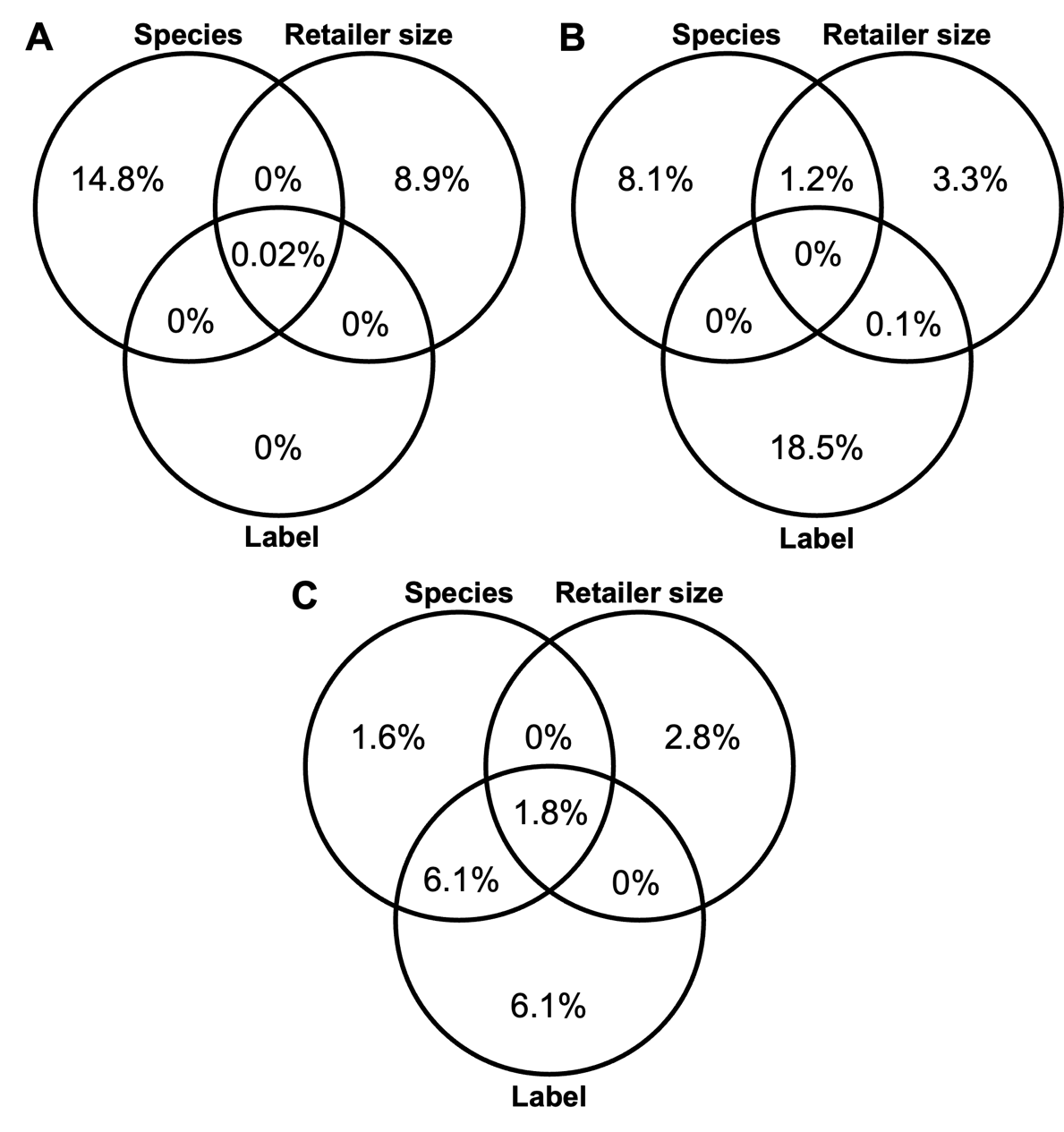

Figure S1. Semi partial variance explained among important predictor variables and sets of important predictor variables in explaining A) richness, B) diversity, and C) exceedances.

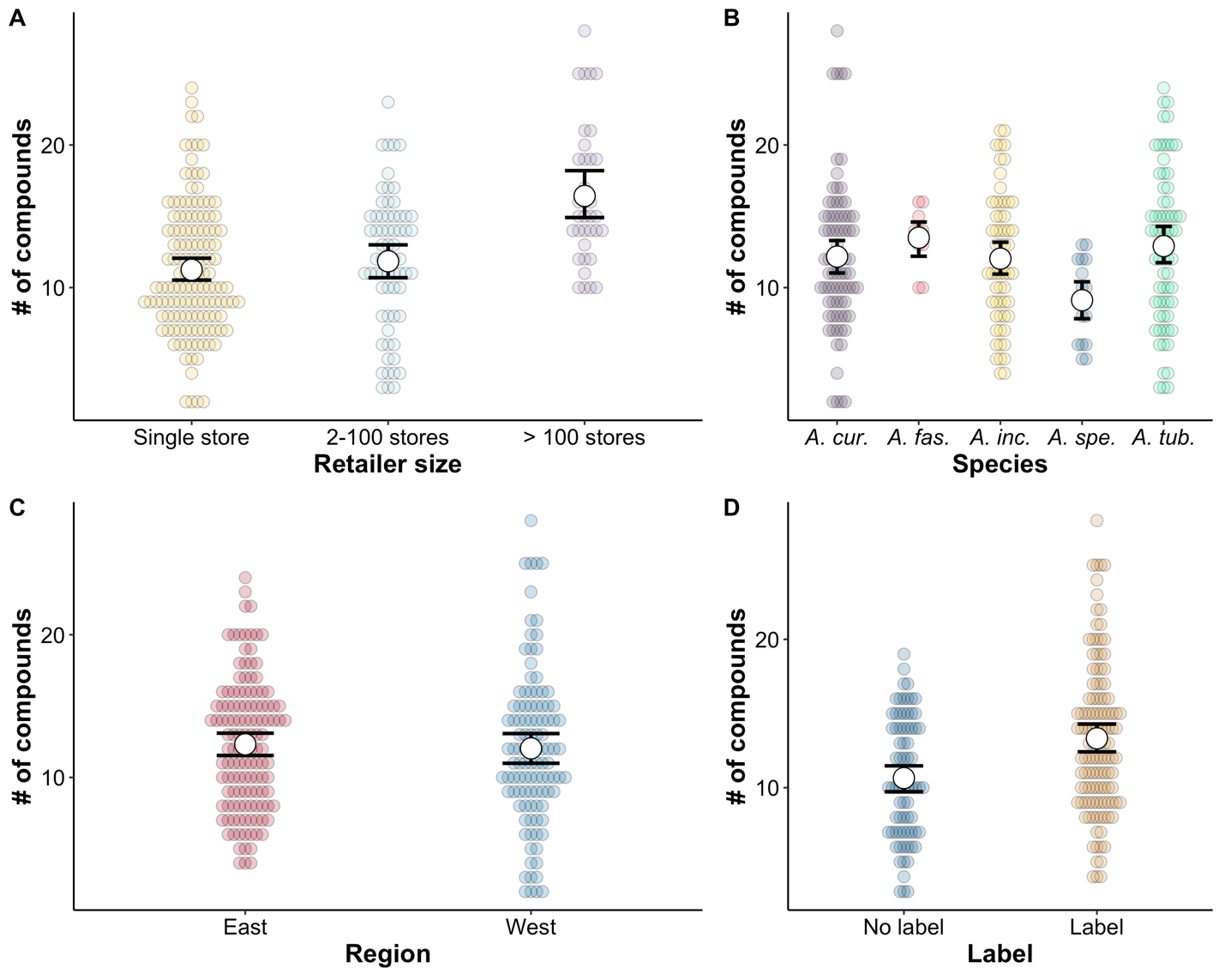

Figure S2. Summary of pesticide richness found in milkweed samples purchased in stores across the United States. Points show the mean effective number of compounds by plant for each variable and the bars show standard errors.

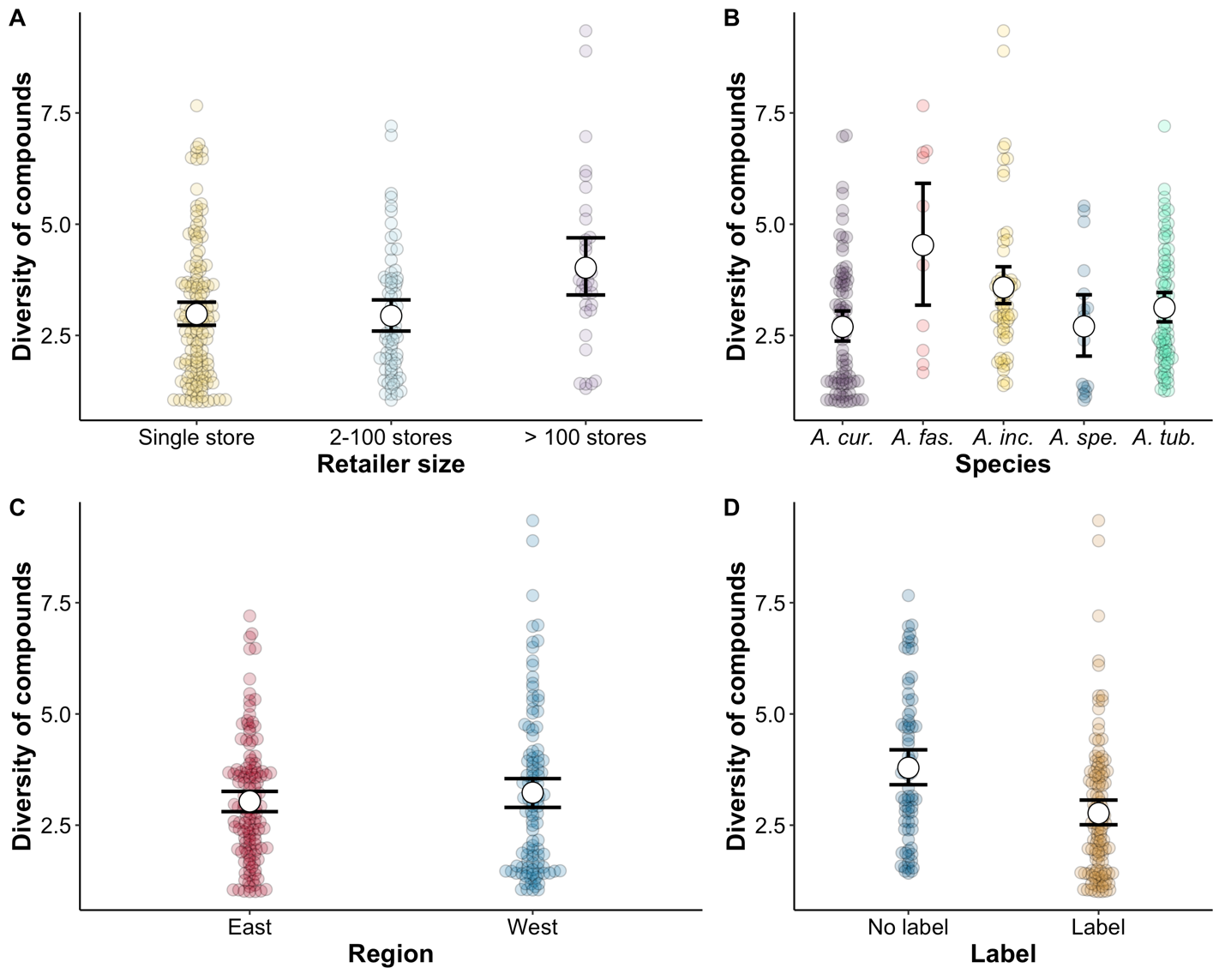

Figure S3. Summary of pesticide diversity found in milkweed samples purchased in stores across the United States. Diversity is represented as the effective number of compounds by taking the exponential of the Shannon diversity index. Points show the mean effective number of compounds by plant for each variable and the bars show standard error.

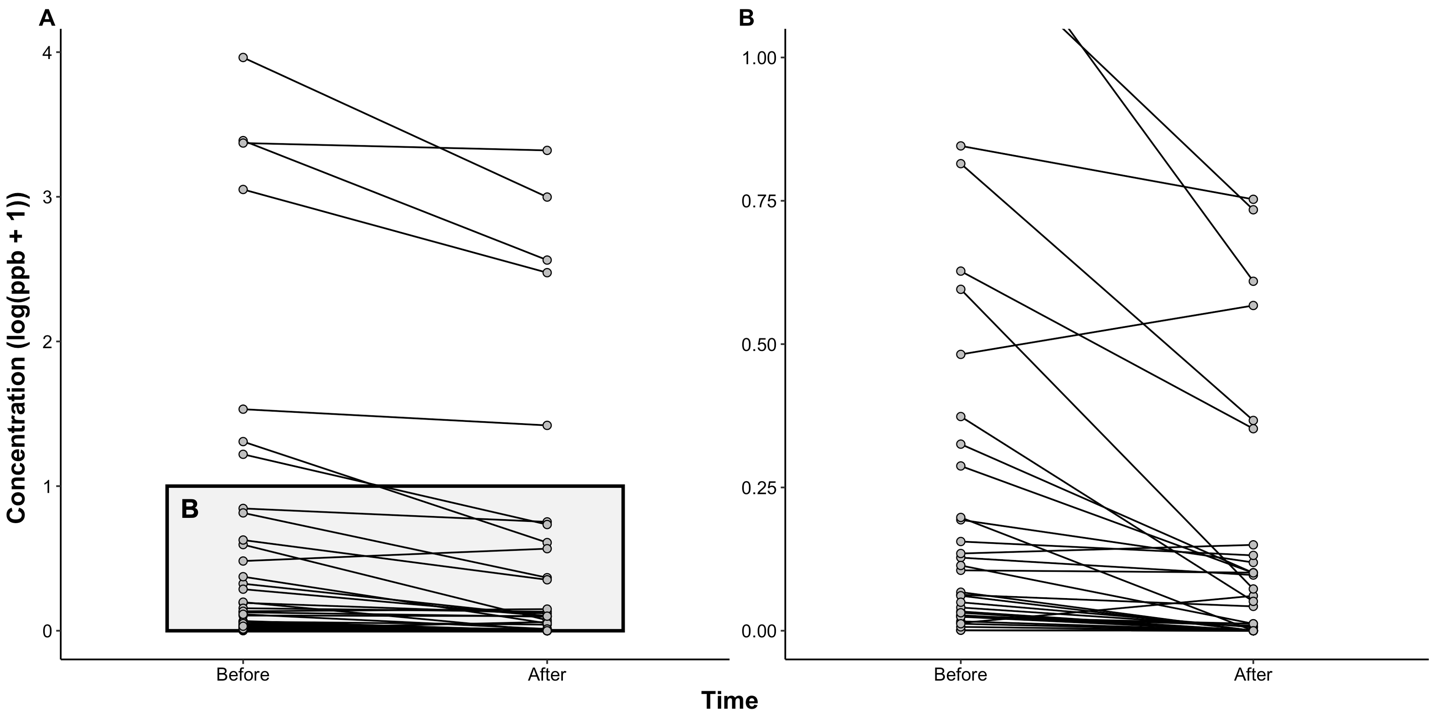

Figure S4. Changes in pesticide concentrations two weeks after purchase. A) The changes in all pesticides and B) changes in low concentration pesticides (the compounds highlighted with the gray box in the left panel).
